# Enhanced cross-modal activation of sensory cortex in mouse models of autism

**DOI:** 10.1101/150839

**Authors:** Ryoma Hattori, Thomas C. Südhof, Kazuhiro Yamakawa, Takao K Hensch

**Affiliations:** Center for Brain Science, Department of Molecular & Cellular Biology, Harvard University, 52 Oxford St, Cambridge, MA 02138 USA; Neurobiology Section, Center for Neural Circuits and Behavior, Department of Neurosciences, University of California, San Diego, La Jolla, CA 92093, USA; Department of Molecular and Cellular Physiology, Stanford University, Stanford,CA 94304-5453, USA; Howard Hughes Medical Institute; Laboratory for Neurogenetics, RIKEN Brain Science Institute, Wako, Saitama, Japan

## Abstract

**SUMMARY:** Synesthesia is a condition wherein one sense is evoked by another. Recent studies suggested a higher incidence of synesthesia among people with autism. However, the underlying circuit mechanism of the comorbidity remains unknown partly due to lack of animal models. Here, we measured auditory response of primary visual cortex (V1) in mouse models to estimate the mixture level of their senses. We found that the V1 auditory response exhibits bidirectional cross-modal plasticity and depends on the level of GABA-mediated inhibition. Analysis of the V1 auditory response in autistic BTBR strain revealed its contralateral bias as in primary auditory cortex, and the auditory evoked field potential was enhanced at gamma range. Furthermore, early sound-driven spike modulation of V1 was commonly shifted toward enhancement in three different autism models (BTBR, NL3 R451C, SCN1A R1407X). Disruption of excitatory/inhibitory (E/I) circuit balance is prevalent among autistic people and mouse models. Thus, our results suggest that E/I imbalance may be the common circuit dysfunction for both autism and synesthesia.

## INTRODUCTION

Synesthesia is a neurological condition in which a certain sense is automatically induced by cross-modal stimuli. The prevalence of synesthesia in general population is estimated as ∼4.4 % (Simner *et al*., 2006), exceeding the prevalence of major mental disorders, for example, ∼1.5 % for autism (Baio *et al*., 2014) and ∼1.1 % for schizophrenia (Regier *et al*. 1993). Nonetheless, the neural and genetic mechanism of synesthesia is much less understood than the others partly due to lack of animal models.

It has been proposed that synesthesia may occur if synaptic pruning of neural connections between different sensory areas fails during development (Bargary & Mitchell, 2008). In line with the pruning hypothesis, anatomical hyper-connectivity between two synesthetically linked sensory areas has been confirmed in some synesthetes (Rouw & Scholte, 2007; Hänggi *et al*., 2008). However, such anatomical hyper-connectivity could be mere a byproduct of reduced inhibition, and disinhibition in sensory area itself might cause synesthesia.

Recent studies reported that synesthesia is more common in people with autism (Baron-Cohen et al., 2013; Neufeld et al., 2013), suggesting a common circuit dysfunction. Although neural connectivity is abnormal in people with autism (Courchesne et al., 2007), GABA-mediated inhibitions are commonly impaired in autistic patients (Coghlan et al., 2012) and its mouse models (Gogolla et al., 2009). In this study, we used auditory response in visual cortex as a measurement of sensory blending in mouse models to explore the candidate mechanism of synesthesia.

## RESULTS

### Auditory response in primary and secondary visual cortex

Although the majority of neurons in visual cortex responds exclusively to visual stimuli, some neurons respond to non-visual sensory stimuli in rodent (Wallace et al., 2004). To visualize the pattern of visual or cross-modal auditory response in visual cortex, we measured intrinsic fluorescent signals from flavoproteins (Tohmi et al., 2006; Gogolla et al., 2014) in response to flash light or sound stimuli. In superficial layer, we detected sound-driven response and audio-visual interaction in both primary (V1) and secondary (V2) visual cortex of adult wildtype (C57Bl6/J) mice (> P60, 23 mice) (Figures 1A-E). When combined with concurrent sound, visual responses in V1 and V2 were further enhanced and sharpened (Figures 1B-1D). Comparison of the multimodality between V1 and V2 using two different indexes (percent enhancement of visual response by sound and auditory response relative to visual response) revealed higher multimodality in V2 (Figure 2E), consistent with an increased multimodality ascending the cortical hierarchy.

**Figure 1.**
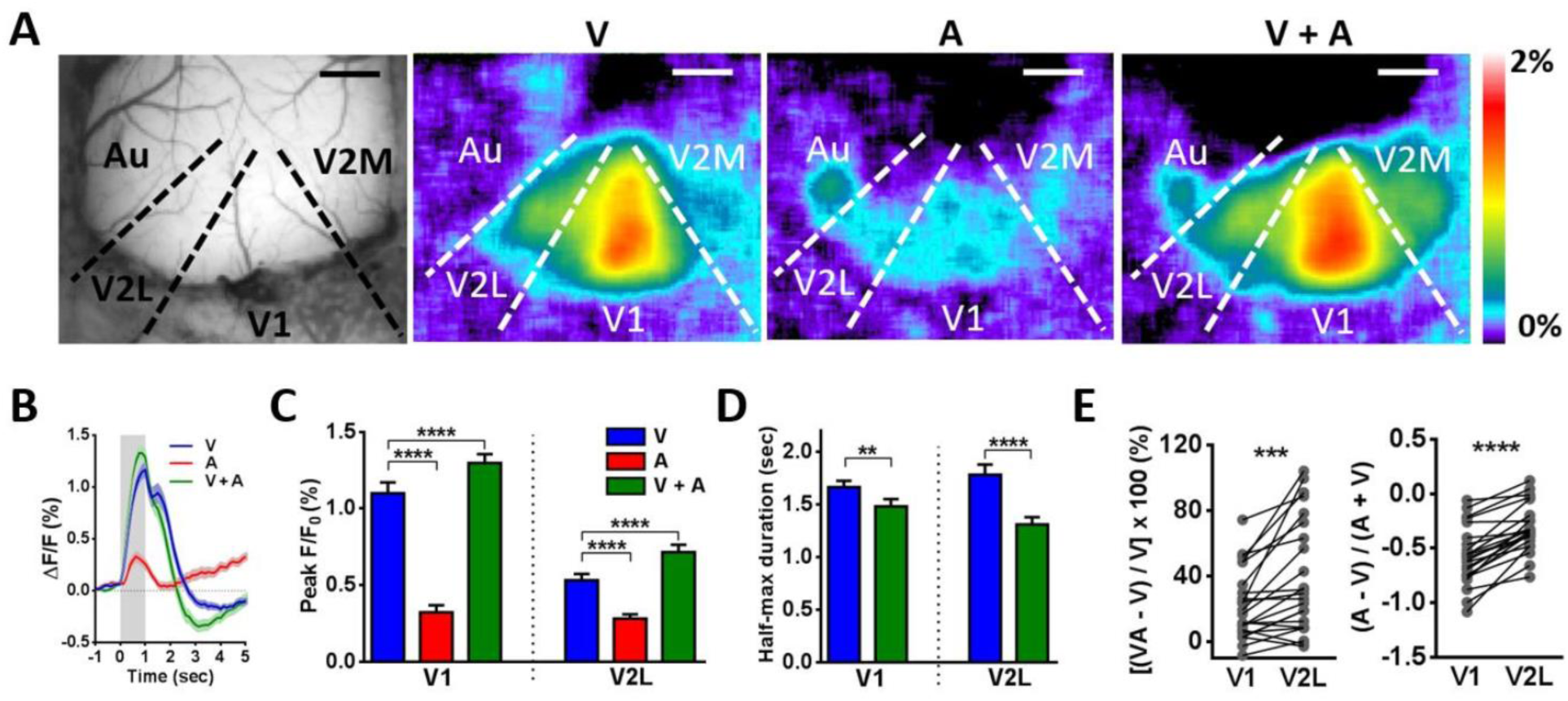
Flavoprotein imaging of visual and cross-modal auditory response in mouse visual cortex. (A) Typical cranial window and activation by visual (V), auditory (A) or combined (V+A) stimuli. Scale bars, 1 mm. V1, primary; V2L, lateral and V2M, medial secondary visual cortex; Au, auditory areas. (B) Average signal traces of flavoprotein signals in V1 (n = 23 mice). (C) Peak amplitudes of flavoprotein signals in V1 and V2L. One-way ANOVA, followed by Holm-Sidak test. (D) Half-maximum durations of flavoprotein signals in V1 and V2L (blue: V, green: combined). Paired t-test. (E) Comparison of multimodality between V1 and V2L with two different indexes. Paired t-test. All data are presented as mean ± SEM. **P < 0.01, ***P < 0.001, ****P < 0.0001.

**Figure 2.**
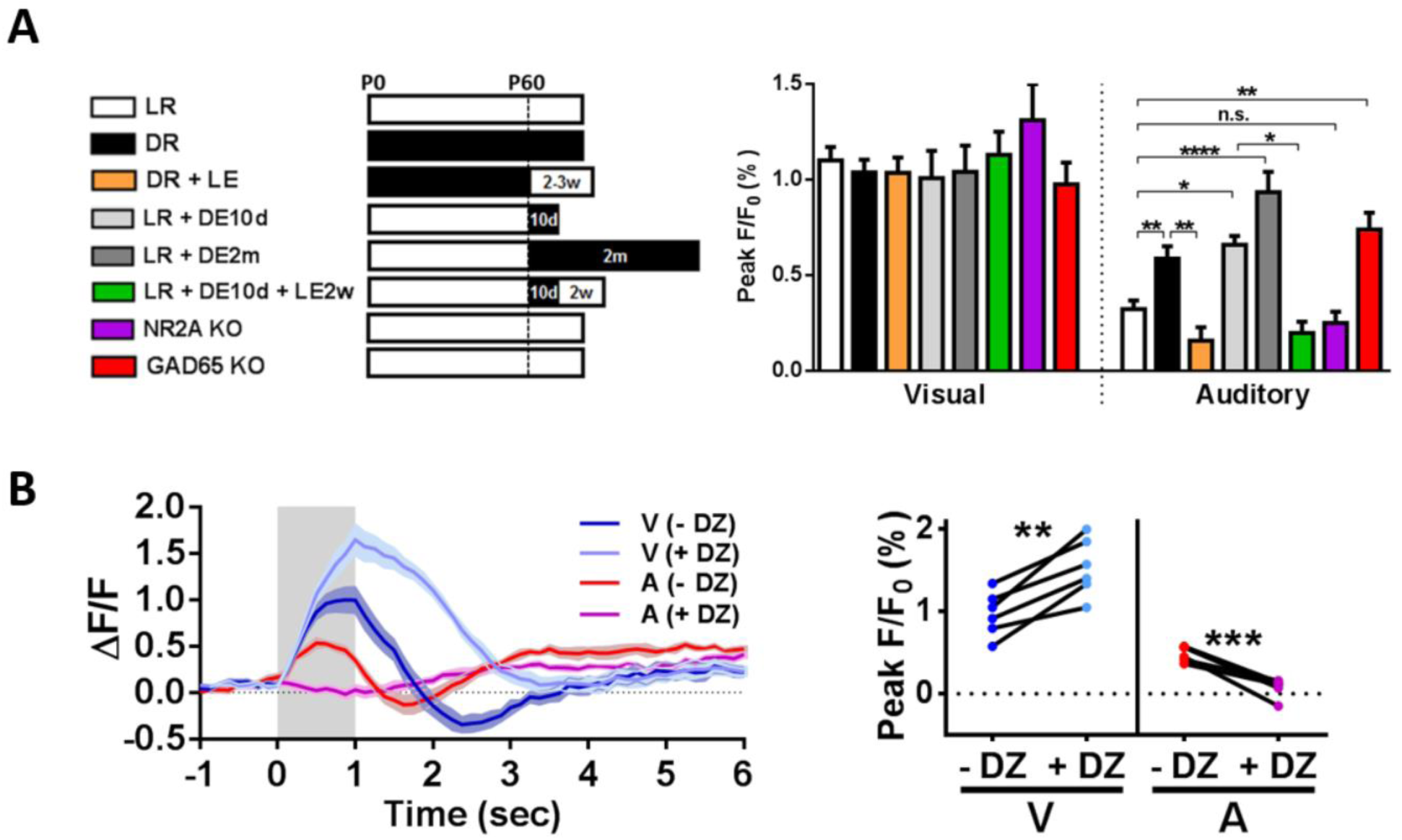
Cross-modal auditory response in V1 is reversibly regulated by visual experience and inhibition. (A) Rearing conditions (white: 12hr light/dark, black: complete darkness) and peak visual or auditory flavoprotein response in V1 (LR: n = 23 mice, DR: n = 16 mice, DR+LE: n = 6 mice, LR+DE10d: n = 5 mice, LR+DE2m: n = 5 mice, LR+DE10d+LE2w: n = 6 mice, NR2A KO: n = 5 mice, GAD65 KO: n = 6 mice). One-way ANOVA, followed by Holm-Sidak test. (B) Average time course of visual or auditory response and their peaks before (-DZ) and after (+ DZ) acute diazepam injection in DR (n = 6 mice). Paired t-test. (C) All data are presented as mean ± SEM. *P < 0.05, **P < 0.01, ***P < 0.001, ****P < 0.0001.

### Darkness enhances cross-modal auditory response

Congenital blindness causes cross-modal plasticity and enhances cross-modal response in human visual cortex (Pascual-Leone et al., 2005). To understand the neural mechanism that regulate cross-modal auditory response in visual cortex, we generated the mouse model of cross-modal plasticity by rearing mice in darkness from birth (DR: dark-reared) until > 2 months of age (16 mice). Consistent with human studies, their cross-modal response was larger than age-matched normally reared controls (LR: light-reared) and returned to LR level after 2-3 weeks of light exposure (6 mice) (Figure 2A).

In adult human visual cortex, such cross-modal plasticity is reversible even with short-term blindfolding or succeeding light exposure (Merabet et al., 2008). To model this adult cross-modal plasticity, we exposed LR mice to darkness for 10 days (5 mice) or 2 months (5 mice) from P60. Both groups of animals robustly enhanced auditory response in their V1 (Figure 2A). Furthermore, enhanced cross-modal response after 10-day dark-exposure (DE) reversibly returned to pre-DE level after re-exposure to light for 2 weeks (6 mice) (Figure 2A).

### Inhibition regulates cross-modal auditory response

Then, what is the neural mechanism that regulates cross-modal auditory response in visual cortex? Given the short time course of adult cross-modal plasticity, formation of new long-range neural connections between auditory and visual cortex is unlikely. In the V1 of adult rodents, 10-day DE dampens cortical inhibitions and increases NR2B/NR2A subunit ratio of NMDA glutamate receptor (He et al., 2006; Huang et al., 2010). In contrast, light-exposure after dark-rearing enhances cortical inhibitions (Morales et al., 2002; Jiang et al., 2010) and decreases NR2B/NR2A ratio (Quinlan et al., 1999).

To examine the contributions of cortical inhibition and NR2B/NR2A ratio, we measured cross-modal auditory response from mice lacking NR2A subunit gene (NR2A KO, 5 mice) (Fagiolini et al., 2003) and mice lacking a synaptic isoform of GABA synthase, GAD65 (GAD65 KO, 6 mice) (Hensch et al., 1998). Although the cross-modal auditory response in NR2A KO mice was comparable to control animals, GAD65 KO mice showed larger cross-modal auditory response without visual deprivation (Figure 2A). Furthermore, acute intraperitoneal injection of diazepam (3 mg/kg, i.p.), a positive allosteric modulator of the GABA_A_ receptor, suppressed cross-modal auditory responses in DR mice (> P60, 6 mice) (Figure 2B). In contrast to cross-modal response, baseline visual responses were not significantly affected by visual deprivation or GAD65 deletion (Figure 2A) but were enhanced by acute diazepam (Figure 2B). Thus, cortical inhibition preferentially suppresses cross-modal response, thereby increasing unimodality in V1.

### Robust cross-modal auditory response in BTBR T+tf/J strain

Impaired GABA-mediated inhibition is prevalent among autistic mouse models (Gogolla et al., 2009), including GAD65 KO mice that exhibit repetitive and asocial behaviors (Gogolla et al., 2014). The heightened prevalence of synesthesia among autistic subjects implies the shared neural mechanism for both symptoms (Baron-Cohen et al., 2013; Neufeld et al., 2013). To test this comorbidity directly, we turned to a well-characterized mouse model of idiopathic autism, the inbred BTBR T+tf/J strain (Silverman et al., 2010), which exhibits weak GABAergic inhibitions (Gogolla et al., 2014; Han et al., 2014).

Indeed, flavoprotein imaging revealed a robust cross-modal auditory response in the V1, which was preferentially evoked by sound coming from their contralateral side (5 mice) (Figure 3A). Contralateral bias of auditory response was also observed in auditory cortex (Figure 3A) as reported previously (Popescu and Polley, 2010).

**Figure 3.**
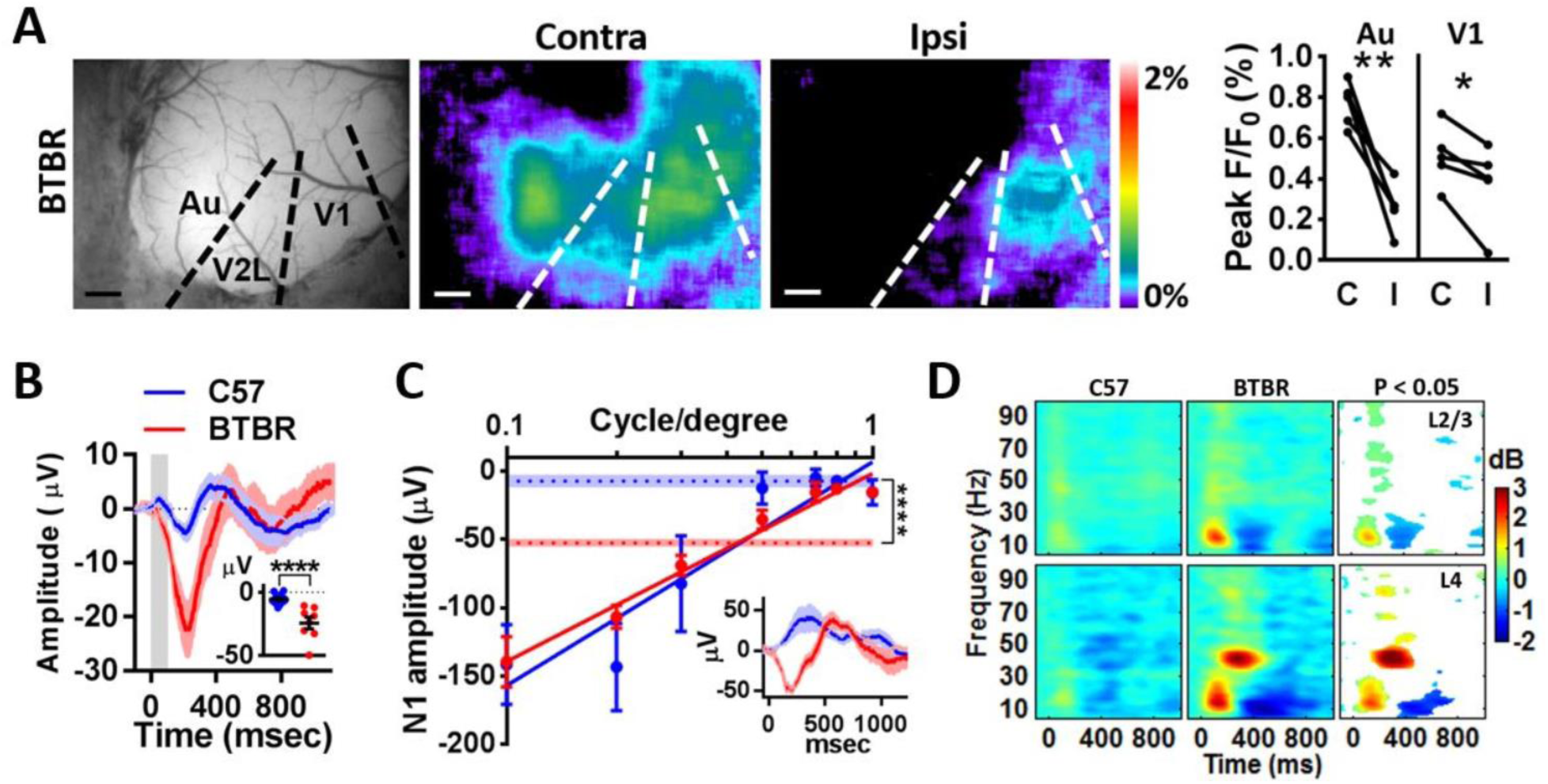
Enhanced cross-modal auditory response in autistic BTBR strain. (A) Cortical responses to white noise sounds from contralateral or ipsilateral side (BTBR; n = 5 mice). Cortical areas demarcated on the basis of LED flash responses. Scale bar, 1 mm. Paired t-test. (B) Average cAEP traces recorded at L2/3 of V1 (C57: n = 16 mice, BTBR: n = 8 mice). Inset compares their first negative peak (N1) amplitudes. Unpaired t-test. (C) N1 amplitudes of VEPs as a function of grating spatial frequency at the L4 of the center of V1 (C57: n = 5 mice, BTBR: n = 4 mice). N1 amplitudes of cAEP (mean ± SEM) at the same recording sites are indicated by dashed lines. Inset shows average cAEP traces at this recording sites. The center of V1 was targeted for each mouse after thinned-skull flavoprotein imaging of visual response. Unpaired t-test. (D) Sound-evoked spectral perturbation from baseline (pre-stimulus baseline of 500 ms). Areas with significant difference between C57 and BTBR at L2/3 or L4 are colored in the right statistics column (pixel-by-pixel unpaired t-test). All data are presented as mean ± SEM. *P < 0.05, **P < 0.01, ****P < 0.0001.

We also directly measured sound-driven electrical activities in V1 by local field potential recordings. Although the negative peak (N1) amplitude of visual-evoked potential (VEP) at the L4 was comparable between C57 strain and BTBR strain (Figure 3C), the N1 amplitude of cross-modal auditory-evoked potential (cAEP) was larger at both L2/3 (C57: 16 mice, BTBR: 8 mice) (Figure 3B) and L4 (C57: 5 mice, BTBR: 4 mice) (Figure 3C) in BTBR strain. Spectral analysis of cAEP revealed a larger sound-driven power enhancement at wide-frequency in BTBR strain, but the most notable difference was at γ frequency range where sound mostly suppressed power in C57 but enhanced power in BTBR (Figure 3D).

### Abnormal audio-visual spike interactions are prevalent among mouse models of autism

To examine the audio-visual interactions at cellular level, we next recorded cross-modal spiking activity from V1. We detected significant sound-driven spiking responses and multisensory spike enhancement in the V1 of BTBR strain with its mean onset latency faster than visual response (241 units; auditory: 12 ms, visual: 30 ms) (Figures 4A and 4B).

**Figure 4.**
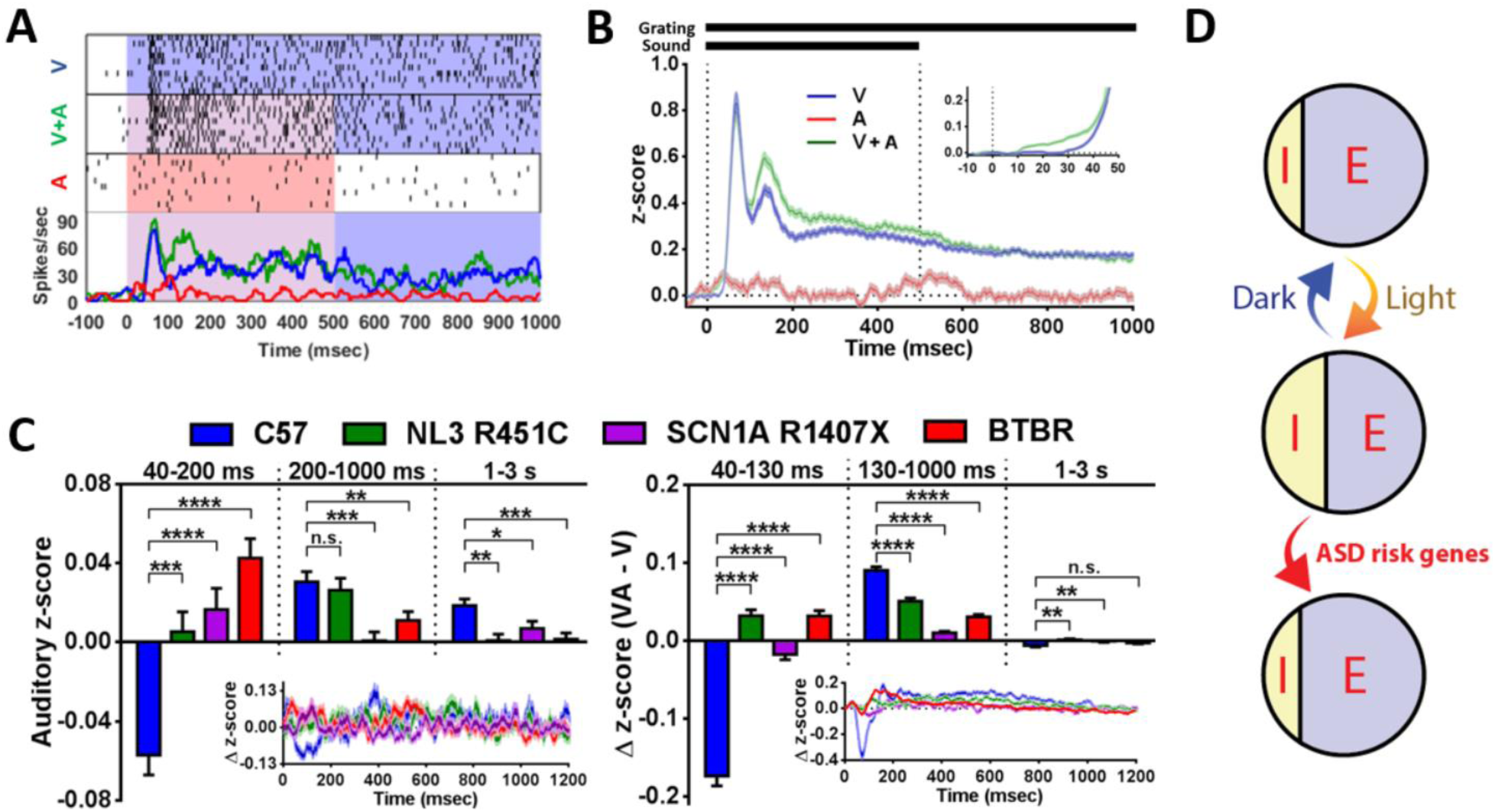
Sound-driven spike modulations of V1 neurons in autistic mouse models. (A) Raster plots and peri-stimulus time histograms (PSTHs) of a typical representative unit from BTBR strain in response to visual (V), auditory (A), and combined (V+A) stimuli. Preferred drifting grating orientation of this unit was used as a visual stimulus for this example. (B) z-scored population averages (mean ± SEM) of PSTHs (> P70; n = 241 units). Inset highlights the pre-visual phase. (C) Auditory z-scores or Δ z-scores between combined (VA) and visual (V) conditions at different post-stimulus time intervals (C57: n = 149 units, NL3 R451C: n = 111 units, SCN1A R1407X : n = 136 units, BTBR: n = 241 units; all mice are > P70). Insets show the z-scores as a function of time (mean ± SEM). One-way ANOVA, followed by Holm-Sidak test. (D) In normally reared C57 strain, sound-driven suppression and enhancement of V1 activities are balanced. Increased excitatory/inhibitory (E/I) ratio in their V1 by either dark-rearing or ASD risk gene mutations shifts the balance of sound-driven V1 modulations toward enhancement. On the other hand, light-exposure of dark-exposed animals re-balances the cross-modal modulations by increasing inhibitions in V1. All data are presented as mean ± SEM. *P < 0.05, **P < 0.01, ***P < 0.001, ****P < 0.0001.

We then compared the sound-driven spike modulations of V1 between non-autistic wild-type C57 strain (149 units) and three different mouse models of autism, BTBR strain, Neuroligin-3 (NL3) R451C knock-in mice (111 units), and Scn1a R1407X knock-in mice (136 units). R451C mutation in NL3 gene is associated with autism in human (Jamain et al., 2003) and causes autistic behaviors in mice (Tabuchi et al., 2007; Burrows et al., 2015). Scn1a is also strongly linked with autism and epilepsy (Zhu et al., 2014), and its loss of function mutations such as R1407X (Ogiwara et al., 2007) result in autistic behaviors in mice (Han et al., 2012; Ito et al., 2013). Although sound transiently suppressed V1 spiking activities in wild-type C57 strain, the sound-driven spike suppression was commonly abolished in all three mouse models of autism (Figures 4C). Consequently, the ratio of sound-driven spike enhancement/suppression was commonly increased in mouse models of autism.

## DISCUSSION

In the present study, we found that visual experience and inhibition critically regulate cross-modal auditory response in mouse V1. Furthermore, comparison of multiple mouse models of autism revealed an impaired sound-driven spike suppression as a common characteristics of autistic mice. Our results indicate that loss of cross-modally driven spike suppression in autism, which likely results from excitatory/inhibitory (E/I) imbalance in autistic brain (Gogolla et al., 2009), unmasks excitatory cross-modal response in primary sensory cortex.

### E/I imbalance as a potential neural mechanism of synesthesia

Enhanced cross-modal auditory response in mouse models of autism suggests synesthesia-like sensory blending in their brain. In humans, synesthesia is either induced by mutations (Brand and Ramachandran, 2011) or acquired by blindfolding or hallucinogen use (Afra et al., 2009). In our study, we provided both genetic models (BTBR, NL3 R451C, Scn1a R1407X) and sensory deprivation models (DR, DE) of increased sensory blending as potential mouse models of synesthesia. E/I is commonly imbalanced in BTBR (Gogolla et al., 2014; Han et al., 2014), NL3 R451C (Cellot and Cherubini, 2014; Speed et al., 2015) and Scn1a mutant (Ogiwara et al., 2007; Tai et al., 2014). Similarly, DR or DE impairs GABA-mediated inhibitions (Morales et al., 2002; He et al., 2006) which is rescued by light-exposure (Morales et al., 2002). Therefore, E/I balance critically gates the cross-modality in sensory cortex (Figure 4D) as it does for autism in prefrontal cortex (Yizhar et al., 2011). Thus, the disinhibited cross-modality may be one of the underlying mechanisms of both autism and synesthesia.

## EXPERIMENTAL PROCEDURES

Please see the Supplemental Experimental Procedures for additional details. All procedures were approved by IACUC of Harvard University or Boston Children’s Hospital.

### Animals

Both sexes were used for each mouse strain. All mice were reared on 12hr light/dark cycle except for dark-reared/dark-exposed mice. For the latter, cages were placed in a light-tight chamber within a dark room.

### Flavoprotein imaging

Mice were first anesthetized with Nembutal (50 mg/kg, i.p.) / Chlorprothixene (8 mg/kg, i.m.), then injected with Atropine (1.5 mg/kg, s.c.). After craniotomy, dura was removed to enhance fluorescence signals. Brain surface was covered with 3% agarose in saline, and a coverslip was placed on top to create a clear window for imaging. Imaging was started ∼1.5hr after initial Nembutal injection, and additional doses (2.5-5 mg/kg, i.p.) were injected through a polyethylene tube (O.D. 0.965 mm) connected to the abdomen whenever necessary to keep animals in a lightly anesthetized state during imaging. Body temperature was maintained at 38°C during imaging. Eyes were covered with silicone oil to prevent drying. For thinned-skull imaging, the skull over V1 was thinned as much as possible before it was covered by 3% agarose and a coverslip.

Each trial lasted 10 sec and was composed of three epochs: pre-stimulation for 3 sec, stimulation for 1 sec, and post-stimulation for 6 sec. After each trial, there was a 10 sec rest period. Three types of stimuli (visual, auditory, audiovisual) were alternately presented, and 50-60 trials were averaged for each modality. Visual stimulus was an LED (λ = 630 nm, luminance = 10 cd, diameter = 5 mm) and the auditory stimulus was a 4.1 kHz piezo buzzer or white noise at ∼80 dB. Both stimuli were placed 10 cm away at 70° azimuth and 35° elevation towards the contralateral eye (peripheral visual field) to distinguish V1 responses from V2L responses (Marshel et al., 2011). For the experiment in Figure 3A, white noise sounds were alternately delivered through plastic tubes inserted into each ear canal to maximize the interaural level difference. For acute Diazepam, a sedative dose (3 mg/kg, i.p.) (Ohl et al., 2001) was injected through a polyethylene tube (O.D. 0.965 mm) connected to the abdomen. Imaging was started 5 min post-injection.

### Field potential recording

Mice were anesthetized with Urethane (0.8 g/kg, i.p.) / Chlorprothixene (8 mg/kg, i.m.), and then injected with Atropine (1.5 mg/kg, s.c.) and Dexamethasone (2.5 mg/kg, s.c.). A craniotomy was made over V1, and the dura was removed before placing a chamber on V1. The chamber was filled with saline. Evoked-potentials were recorded at a depth of ∼250 μm for L2/3 and ∼400 μm for L4. Body temperature was maintained at 38°C, and O_2_ gas was continuously provided through a trachea tube. Both eyes were sutured shut during cAEP recordings to prevent potential visual inputs. White noise bursts (80 dB, 100 msec duration) were repeatedly presented every 2.5 sec from a speaker placed contralaterally, and ∼500 trials were averaged for each animal. For VEP recordings, the ipsilateral eye was kept closed by lid suture, and the exposed contralateral eye was covered by silicone oil. Flickering horizontal gratings with various spatial frequencies were presented every 1.25 sec alternately with or without white noise sound bursts, and 400 trials of each condition were averaged separately.

### Single-unit recordings

Mice were anesthetized with Urethane (0.8 g/kg, i.p.) / Chlorprothixene (8 mg/kg, i.m.), and then injected with Atropine (1.5 mg/kg, s.c.) and Dexamethasone (2.5 mg/kg, s.c.). A craniotomy was made over V1, but the dura was kept intact. Spikes were recorded (Plexon) using a linear silicon probe with 16 channels (NeuroNexus Technologies). The brain was covered by 3% agarose in saline. Body temperature was maintained at 37°C, and O_2_ gas continuously provided through a trachea tube. Drifting gratings (0.03 cycle/degree, 3 sec duration followed by 3 sec grey screen) with 24 different orientations (360°/15°) were presented in randomized order alternately with or without a white noise burst (80 dB, 500 msec duration, concurrent onset with grating) from an electrostatic speaker (Tucker-Davis Technologies) placed next to contralateral ear. Ipsilateral eye was covered by a piece of black tape to limit visual inputs to contralateral eye. ∼10 trials were averaged for each condition. Sound effects on visual spiking activities evoked by 24 orientations were further averaged to quantify audio-visual interaction. After recordings, spike waveforms were sorted by Offline Sorter (Plexon) and data analysis was performed in MATLAB.

## ACKNOWLEDGEMENTS

We thank M Nakamura, E Centofante and N. De Souza for mouse colony maintenance; D Brady, N Gogolla and N Picard for advice on flavoprotein imaging and single-unit recording, respectively. Funded by NIMH Silvio Conte Center (1P50MH094271 to TKH) and the Nakajima Foundation (RH).

## AUTHOR CONTRIBUTIONS

R.H. designed the study, performed the experiments, interpreted the results, and wrote the manuscript. T.S.C generated and provided NL3 R451C mice. K.Y. generated and provided SCN1A R1407X mice. T.K.H. supervised the study.

## Supplemental Information

## SUPPLEMENTAL EXPERIMENTAL PROCEDURES

### Animals

Founders of C57BL6/J and BTBR T+tf/J inbred mouse strains were purchased from Jackson laboratory. GAD65 KO, NR2A KO and NL3 R451C KI were originally generated by Drs. K. Obata (Asada et al., 1996), M. Mishina (Sakimura et al., 1995), T. C. Südhof (Tabuchi et al., 2007), respectively, and backcrossed to C57BL6/J mice. Both sexes were used. All mice were reared on 12hr light/dark cycle except for dark-reared/dark-exposed mice. For the latter, cages were placed in a light-tight chamber within a dark room. All procedures were approved by IACUC of Harvard University or Boston Children’s Hospital.

### Flavoprotein imaging

Mice were first anesthetized with Nembutal (50 mg/kg, i.p.) / Chlorprothixene (8 mg/kg, i.m.), then injected with Atropine (1.5 mg/kg, s.c.). A craniotomy was made to expose V1 / V2, and the dura was carefully removed to enhance the auto-fluorescence signal. The brain surface was covered with 3% agarose in saline, and a coverslip was placed on top to create a clear window for imaging. Gentle pressure was applied to the brain tissue to flatten visual cortex when the coverslip was placed. Any exposed agarose was covered with silicone oil and plastic wrap to prevent drying. Imaging was started ∼1.5hr after initial Nembutal injection, and additional doses (2.5-5 mg/kg, i.p.) were injected through a polyethylene tube (O.D. 0.965 mm) connected to the abdomen whenever necessary to keep animals in a lightly anesthetized state during imaging.

Body temperature was maintained at 38°C during imaging. Eyes were covered with silicone oil to prevent drying. For thinned-skull imaging, the skull over V1 was thinned as much as possible before it was covered by 3% agarose and a coverslip.

Blue light (470-490 nm) was used as excitation and the autofluorescence (500-550 nm) from flavoproteins was recorded (8 frames/sec) with a cooled CCD camera on a dissecting microscope (Leica MZ6). After finding the best focal plane with maximal cross-modal auditory response, autofluorescence in response to sensory stimulation was acquired using Metamorph (Molecular Devices). Each trial lasted 10 sec and was composed of three epochs: pre-stimulation for 3 sec, stimulation for 1 sec, and post-stimulation for 6 sec. After each trial, there was a 10 sec rest period. Three types of stimuli (visual, auditory, audiovisual) were alternately presented, and 50-60 trials were averaged for each modality. Visual stimulus was an LED (λ = 630 nm, luminance = 10 cd, diameter = 5 mm) and the auditory stimulus was a 4.1 kHz piezo buzzer or white noise at ∼80 dB. Both stimuli were placed 10 cm away at 70° azimuth and 35° elevation towards the contralateral eye (peripheral visual field) to distinguish V1 responses from V2L responses (Marshel et al., 2011). For the experiment in Figure 3A, white noise sounds were alternately delivered through plastic tubes inserted into each ear canal to maximize the interaural level difference.

For acute Diazepam, a sedative dose (3 mg/kg, i.p.) (Ohl et al., 2001) was injected through a polyethylene tube (O.D. 0.965 mm) connected to the abdomen. Imaging was started 5 min post-injection.

Analysis of images was carried out in Metamorph and MATLAB. The 50-60 trials for each modality were averaged and normalized to the pre-stimulus period (first 20 frames). ROI circles (∼0.75 mm diameter) were placed on the peak visual response in V1 and V2L. Additional reference ROIs were placed outside the craniotomy. The time course of fluorescence change in each ROI was calculated by subtracting signal intensity of the reference ROI at each time point. Peak amplitudes in bar graphs represent averages ± 0.5 sec around each peak. Response pattern images (Figures 1A, 3A) were created by projecting maximum signal intensity in each pixel across entire frames of low-pass (10 x 10 pixels) square-filtered images to a single frame.

### Field potential recording

Mice were anesthetized with Urethane (0.8 g/kg, i.p.) / Chlorprothixene (8 mg/kg, i.m.), and then injected with Atropine (1.5 mg/kg, s.c.) and Dexamethasone (2.5 mg/kg, s.c.). A craniotomy was made over V1, and the dura was removed before placing a chamber on V1. The chamber was filled with saline. A tungsten electrode (∼10 MΩ, FST) was inserted perpendicularly into V1 (∼2.5 mm from lambda), and the signal was band-pass filtered (1-100 Hz). Evoked-potentials were recorded at a depth of ∼250 μm for L2/3 and ∼400 μm for L4. Body temperature was maintained at 38°C, and O_2_ gas was continuously provided through a trachea tube. Both eyes were sutured shut during cAEP recordings to prevent potential visual inputs. White noise bursts (80 dB, 100 msec duration) were repeatedly presented every 2.5 sec from a speaker placed contralaterally, and ∼500 trials were averaged for each animal. For VEP recordings, the ipsilateral eye was kept closed by lid suture, and the exposed contralateral eye was covered by silicone oil. Flickering horizontal gratings with various spatial frequencies were presented every 1.25 sec alternately with or without white noise sound bursts, and 400 trials of each condition were averaged separately.

Data analysis was carried out using MATLAB. Evoked potentials were normalized to their pre-stimulus period (-200 ms to 0 ms). EEGLAB toolbox (Delorme and Makeig, 2004) was used to compute sound-driven spectral perturbations from baseline. Spectral perturbations were calculated using Morlet wavelets and normalized to their pre-stimulus periods (-500 ms to 0 ms).

### Single-unit recordings

To draw PSTHs, firing rates were calculated using 25 ms bins with their centers 2 ms apart. Raw PSTHs were then converted to z-scores by 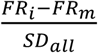 where *FR*_*i*_ is the firing rate in the i^th^ bin of the PSTH, *FR*_*m*_ is the mean firing rate over the entire recording period, and *SD*_*all*_ is the standard deviation of firing rates for the entire recording period. Each z-score was further normalized to baseline by subtracting the mean baseline z-score value of each cell. Mean onset latency was defined as the time when the population averaged PSTH first exceeded the mean + 2SD of baseline (pre-stimulus period: -1000 to 0 ms).

